# Increased ParB level affects expression of stress response, adaptation and virulence operons and potentiates repression of promoters adjacent to the high affinity binding sites *parS3* and *parS4* in *Pseudomonas aeruginosa*

**DOI:** 10.1101/151340

**Authors:** Adam Kawałek, Krzysztof Głąbski, Aneta Agnieszka Bartosik, Anna Fogtman, Grażyna Jagura-Burdzy

## Abstract

Similarly to its homologs in other bacteria, *Pseudomonas aeruginosa* partitioning protein ParB facilitates segregation of newly replicated chromosomes. Lack of ParB is not lethal but results in increased frequency of anucleate cells production, longer division time, cell elongation, altered colony morphology and defective swarming and swimming motility. Unlike in other bacteria, inactivation of *parB* leads to major changes of the transcriptome, suggesting that, directly or indirectly, ParB plays a role in regulation of gene expression in this organism.

ParB overproduction affects growth rate, cell division and motility in a similar way as ParB deficiency. To identify primary ParB targets, here we analysed the impact of a slight increase in ParB level on *P. aeruginosa* transcriptome. ParB excess, which does not cause changes in growth rate and chromosome segregation, significantly alters the expression of 176 loci. Most notably, the mRNA level of genes adjacent to high affinity ParB binding sites *parS1*-*4* close to *oriC* is reduced. Conversely, in cells lacking either *parB* or functional *parS* sequences the orfs adjacent to *parS3* and *parS4* are upregulated, indicating that direct ParB-*parS3/parS4* interactions repress the transcription in this region. In addition, increased ParB level brings about repression or activation of numerous genes including several transcriptional regulators involved in SOS response, virulence and adaptation. Overall, our data support the role of partitioning protein ParB as a transcriptional regulator in *Pseudomonas aeruginosa*.

## Introduction

Accurate copying and segregation of genetic material to progeny cells is crucial for survival and maintenance of species identity. The process of bacterial DNA segregation was first deciphered for low-copy-number plasmids [1–4]. Active partitioning of such plasmids during cell division requires three specific plasmid-encoded factors: an NTPase (A-component), a DNA-binding protein (DBP, B-component) and a special site in DNA, designated centromere-like sequence (*parS*/*parC*). Interactions of DBP with *parS* sequence(s) lead to formation of segrosomes which are then separated by a dynamic NTPase machinery to polar positions assuring their proper segregation during the subsequent cell division [2,3]. Plasmidic active partition systems have been classified into three groups based on the type of NTPase and structure of DBP [2,4]. Homologs of plasmidic Type IA partition proteins, Walker-type ATPases (ParAs) and large DBPs with helix-turn-helix motifs (ParBs), which after binding to *parS* spread on DNA and form large nucleoprotein complexes [5], are also encoded on the majority of bacterial chromosomes [3,4,6]. Multiple copies of highly conserved *parS* sequences are mainly clustered in the so-called *ori* domain comprising ca. 20% of the chromosome [7].

The role of the ParABS systems in accurate bacterial chromosome segregation is widely acknowledged but varies from essential, as exemplified by *Caulobacter crescentus* [8] or *Myxococcus xanthus* [9], to accessory, as in *Bacillus subtilis* [10–13], *Streptomyces coelicolor* [14]*, Vibrio cholerae* [15,16] or *Pseudomonas aeruginosa* [17–19]. Apart from their well-established role in the segregation of newly replicated *ori* domains through DNA compaction [20–22], proper positioning of *ori* domains in the cell [19,23], Par proteins have also been shown to play a role in the control of DnaA activity and replication initiation [24,25] as well as in coordination of cell cycle and differentiation [26–28]. Chromosomal ParB homologs bind to *parS in vitro*, polymerize on DNA and bridge distant sequences [13,29–32]. Whole-genome analyses using chromatin immunoprecipitation (ChIP) have demonstrated spreading of ParB homologs around *parS* sites for up to 20 kb [5,26–29]. Despite the ParB spreading, a transcriptomic analysis in *B. subtilis* did not identify any significant changes in gene expression in a *spo0J* null mutant (Spo0J is a ParB homolog) relative to a WT strain [35]. Limited ParB-dependent transcriptional silencing in the proximity of *parS* sequences has been observed only for several genes in *V. cholerae* [34] and *S. pneumoniae* [36].

In *P. aeruginosa* a lack of ParA and/or ParB is not lethal but results in up to 1000-fold increased frequency of production of anucleate cells even during growth in optimal conditions [17,18,37]. Various *par* mutants exhibit longer division time, increase in cell size, altered colony morphology and are impaired in swarming and swimming motility [17,18]. Ten *parS* sites scattered in the chromosome of *P. aeruginosa* have been identified, but only four of them closest to *oriC* seem to be involved in chromosome segregation [33,38]. A transcriptomic analysis of *P. aeruginosa parA* and *parB* mutants has demonstrated changes in expression of hundreds of loci [39], including genes related to stress response but also many known and putative transcriptional regulators, suggesting a direct and/or indirect role of Par proteins in the regulation of gene expression. In test plasmids, ParB of *P. aeruginosa* was found to spread around *parS* and silence nearby promoters [37], but a comparison of the *parB* mutant and WT transcriptomes did not reveal any obvious changes in the expression of genes adjacent to chromosomal *parS* sequences [39]. However, a recent ChIP-seq analysis of ParB distribution on the *P. aeruginosa* chromosome has revealed, in addition to the region around the high affinity *parS1-4* sequences, also secondary ParB binding sites, apparently not related to known *parS* sites [33].

Similarly to the lack of ParB, also its excess in *P. aeruginosa* affects the cell cycle, as highlighted by slower growth rate, and causes cell elongation and defects in swarming and swimming. ParB excess manifests in nucleoid condensation and increased frequency of anucleate cells formation [17,37,40,41]. The toxicity of ParB overproduction is significantly diminished when ParB is impaired in its polymerization domain [40] whereas ParB variants defective in interactions with DNA, either due to the inability to form dimers or mutations in the DNA binding domain, demonstrate no toxicity [40,41]. This suggests that DNA binding and spreading of ParB protein is crucial for its biological function.

In this project we analysed changes in the transcriptome of *P. aeruginosa* cells with a slightly increased ParB level, not affecting growth rate and chromosome segregation, to define the primary targets of ParB.

## Materials and methods

### Bacterial strains and growth conditions

Bacterial strains used in this study are listed in Table 1. Cultures were grown in L broth [42] or on L agar (L broth solidified with 1.5% agar) at 37°C. For selection of plasmids *Escherichia coli* media were supplemented with 150 μg ml^-1^ (liquid cultures) or 300 μg ml^-1^ (solid media) penicillin (Pn), 10 μg ml^-1^ chloramphenicol (Cm) or 50 μg ml^-1^ kanamycin (Km). Chloramphenicol at concentration 75 μg ml^-1^ (liquid cultures) and 150 μg ml^-1^ (solid media) was used to maintain plasmids in *P. aeruginosa*. 300 μg ml^-1^ carbenicillin (Cb) and 300 μg ml^-1^ rifampicin (Rif) was added to the solid medium for the selection of *P. aeruginosa* transconjugants in allele exchange procedure. To induce the expression from the *araBAD* promoter arabinose (Ara) was added to 0.02% to cultures of PAO1161 with pKGB8 (*araBAD*p) or pKGB9 (*araBAD*p-*parB*) and to 0.1% in PAO1161::*araBAD*p or PAO1161::*araBAD*p-*flag-parB* cultures. For MIC analysis of PAO1161 (pKGB8) and PAO1161 (pKGB9) and PAO1161 as a control, strains were grown in Mueller Hinton cation adjusted medium in the range of Ara concentrations (0, 0.02 and 0.2%) and in the presence of a gradient of piperacillin, ticarcillin, ciprofloxacin, gentamicin, tobramycin or imipenem.

**Table 1.** Bacterial strains used in this study.

| Strain | Description | Reference |
| --- | --- | --- |
| *Escherichia coli* |  |  |
| DH5α | F<sup>-</sup> Φ80lacZΔM15 $\Delta$(lacZYA-argF) U169 recA1 endA1<br>hsdR17 (r<sub>K</sub><sup>-</sup>, m<sub>K</sub><sup>+</sup>) phoA supE44 $\lambda$ thi-1 gyrA96 relA1 | [43] |
| S17-1 | pro ΔhsdR hsdM<sup>+</sup> recA Tp<sup>R</sup> Sm<sup>R</sup> ΩRP4-Tc::Mu Kn::Tn7 | [44] |
| *Pseudomonas aeruginosa* |  |  |
| PAO1161 | PAO1161 Rif<sup>R</sup> leu | [17] |
| PAO1161 parB<sub>null</sub> | PAO1161 Rif<sup>R</sup> leu parB1-18::tetM, Tc<sup>R</sup> | [17] |
| PAO1161 parS<sub>null</sub> | PAO1161 Rif<sup>R</sup> with ten parS sequences mutated or deleted to impair ParB binding to these sequences | [38] |
| PAO1161::araBADp | PAO1161 Rif<sup>R</sup> with araC araBADp T<sub>rrnB</sub><sup>1</sup> cassette integrated in intergenic region between PA5412 and PA5413 (at position 6091513 nt) | this study |
| PAO1161::araBADp-flag-parB | PAO1161 Rif<sup>R</sup> with araC araBADp-flag-parB T<sub>rrnB</sub> cassette integrated in intergenic region between PA5412 and PA5413 (at position 6091513 nt) | this study |
| PAO1161::bexRp-lacZ | PAO1161 Rif<sup>R</sup> Cb<sup>R</sup> with bexRp-lacZ cassette integrated in intergenic region between PA5412 and PA5413 (at position 6091513 nt) | this study |

<sup>1</sup>T<sub>rrnB</sub> transcriptional terminator of *E. coli* rrnB gene

### Transformation and conjugation procedures

Transformation of *E. coli* and *P. aeruginosa* was performed by standard procedures [43,45]. Conjugation between transformants of *E. coli* S17-1 and *P. aeruginosa* PAO1161 Rif^R^ was performed on solid media as described previously [37].

### Plasmids and DNA manipulations

Plasmids used in this study are listed in Table 2. Plasmid manipulations were carried out by standard procedures [46]. Oligonucleotides used in this work are listed in Table S1. All new plasmid constructs were verified by sequencing at the Laboratory of DNA Sequencing and Oligonucleotide Synthesis, IBB PAS.

**Table 2.**
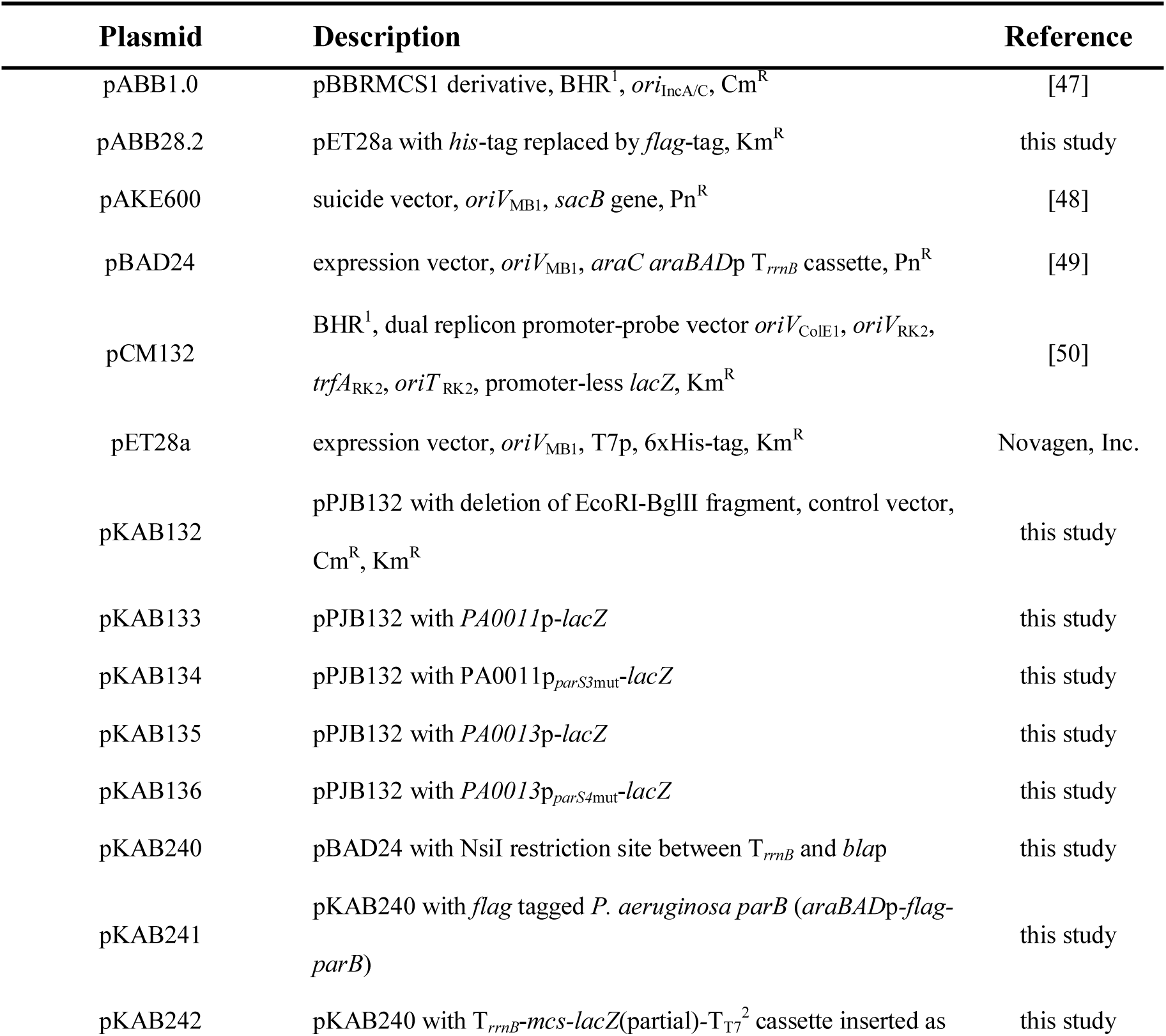
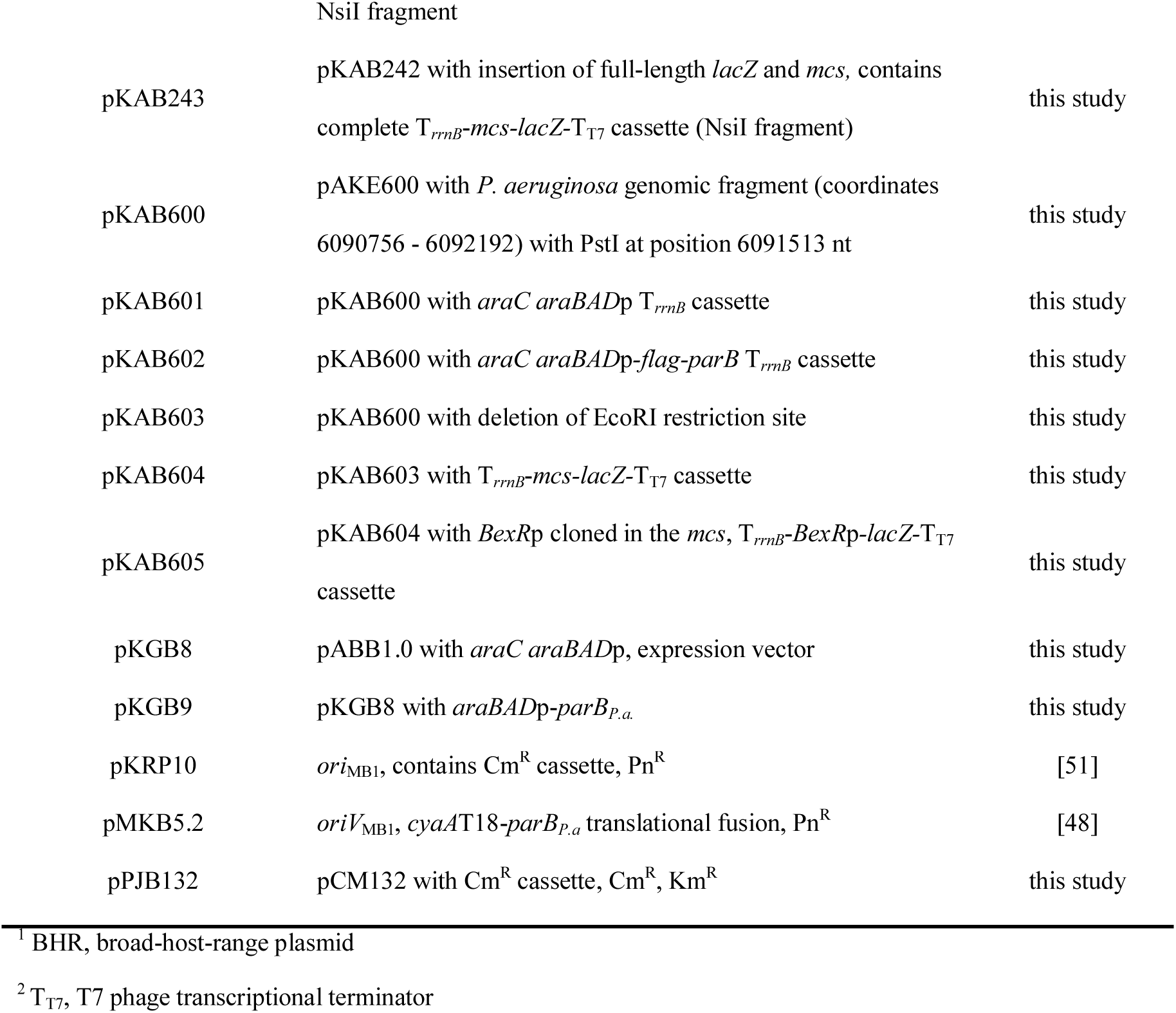
Plasmids used in this study.

Plasmid pKGB8 was constructed by ligation of Eco47III-PstI fragment from pBAD24 containing *araC araBAD*p with HincII and PstI digested pABB1.0. Subsequently, EcoRI-SacI fragment from pMKB5.2 containing *parB* ORF was cloned into pKGB8 to yield plasmid pKGB9.

pET28a (Novagen, Inc.) was modified to add *flag*-tag instead of *his*-tag to the coding sequences. Oligonucleotides #1 and #2 were combined, heated to 100°C for 5 minutes and left for annealing at room temperature. pET28a vector was digested using NcoI and NdeI and ligated with obtained synthetic DNA fragment to yield pABB28.2.

To facilitate transfer of the *araC-araBAD*p-T*_rrnB_* cassette, an additional NsiI restriction site was introduced into pBAD24 downstream of T*_rrnB_*. Two fragments amplified by PCR using pBAD24 as a template and primer pairs #3/#4 and #5/#6, respectively, were used as a template in the second round of PCR with primers #3 and #6 to obtain a 1160 bp product that replaced the HindIII-ScaI fragment in pBAD24 to yield pKAB240.

The *flag*-*parB* fusion was constructed using similar strategy. The *flag* sequence amplified from pABB28.2 using primers #7/#8 and *P. aeruginosa parB* gene amplified from PAO1161 genomic DNA using primers #9/#10 were combined and used as a template in overlap PCR with primers #7/#10. The obtained PCR fragment with *flag-parB* fusion was digested with EcoRI and SalI and ligated into pKAB240 to yield pKAB241.

To construct PAO1161::*araBAD*p and PAO1161::*araBAD*p-*flag*-*parB* strains, the suicide plasmids pKAB601 and pKAB602 were constructed. First, the intergenic region between *PA5412* and *PA5413* from PAO1161 chromosome was amplified in two parts with pairs of primers #11/#12 and #13/#14. Primers #12 and #13 introduced a PstI site. The obtained PCR fragments were digested either with MfeI and PstI or PstI and BamHI, respectively, and the mixture was ligated with EcoRI and BamHI digested pAKE600 [48] to yield pKAB600. The *araC*-*araBAD*p-T*_rrnB_* cassette from pKAB240 and *araC*-*araBAD*p-*flag-parB*-T*_rrnB_* cassette from pKAB241 were excised as NsiI fragments and ligated into PstI site of pKAB600 to yield pKAB601 and pKAB602, respectively. *E. coli* S17-1 strain was transformed with pKAB601 or pKAB602 and the transformants were used as donor strains in conjugation with PAO1161 Rif^R^ [18]. The cassettes were integrated in the defined region of PAO1161 using the allele exchange procedure [18] to obtain PAO1161::*araBAD*p and PAO1161::*araBAD*p-*flag*-*parB*.

pCM132 was modified by insertion of Cm^R^ cassette (SphI fragment from pKRP10 [51]) into SphI site, downstream of *lacZ*, to yield pPJB132. Putative promoter sequences preceding *PA0011* and *PA0013* were amplified on PAO1161 genomic DNA using primer pairs #15/#16 and #17/#18, respectively. The *PA0011*p*_parS3_*_mut_ and *PA0013*p*_parS4_*_mut_ fragments were amplified on PAO1161 *parS*_null_ genomic DNA [38] as a template using the same primer pairs. PCR products were digested with EcoRI and BamHI and ligated between EcoRI and BglII sites into pPJB132 to yield plasmids pKAB133 (*PA0011*p-*lacZ*), pKAB134 (*PA0011*p*_parS3_*_mut_-*lacZ*), pKAB135 (*PA0013*p-*lacZ*) and pKAB136 (*PA0013*p*_parS4_*_mut_-*lacZ*). The empty control vector for β-galactosidase measurements (pKAB132) was obtained by digestion of pPJB132 with EcoRI and BglII, blunting by fill-in with Klenow fragment of DNA PolI and self-ligation.

To construct PAO1161 Rif^R^ strain with *bexR*p-*lacZ* cassette integrated in intergenic region between *PA5412* and *PA5413*, the suicide plasmid pKAB605 was constructed. First, the T*_rrnb_* with *mcs*, part of the *lacZ* gene and T_T7_ was amplified using overlap PCR strategy. To this end, 2 fragments were amplified on pCM132 plasmid as a template using primer pairs #19/#20 and #21/#22, respectively, and the 3rd fragment was amplified on pET28a using primer pair #23/#24. The three fragments were mixed and used as a template in an overlap PCR with primers #19/#24. The obtained 787 bp product digested with NsiI replaced the *araC*-*araBAD*p-T*_rrnB_* in pKAB240 yielding plasmid pKAB242. *mcs*-*lacZ* was subsequently excised from pCM132 as EcoRI and BlpI fragment and ligated with EcoRI-BlpI digested pKAB242 to yield pKAB243, containing complete T*_rrnB_*-*mcs-lacZ-*T_T7_ cassette.

The single EcoRI restriction site in pKAB600 plasmid was removed by EcoRI digestion, filling in by Klenow fragment of DNA PolI and plasmid self-ligation to yield pKAB603. The T*_rrnB_*-*mcs-lacZ-*T_T7_ cassette excised from pKAB243 by NsiI digestion was ligated into PstI site of pKAB603 to yield pKAB604. The region preceding *bexR* (*PA2432*) gene was amplified on PAO1161 genomic DNA using primers #25/#26, digested with EcoRI and BamHI and ligated with EcoRI and BglII digested pKAB604 to yield plasmid pKAB605 carrying T*_rrnB_*- *bexR*p*-lacZ-*T_T7_ cassette flanked by sequences allowing integration in the intergenic region between *PA5412* and *PA5413*. *E. coli* S17-1 strain transformed with pKAB605 was used as donor in conjugation with PAO1161 Rif^R^ [18]. The suicide vector was integrated in the defined region of PAO1161 (at position 6091513 nt) using homology recombination [18] and the carbenicillin-resistant integrant (PAO1161::*bexR*p-*lacZ*) was used as a recipient in a subsequent conjugation with S17-1 cells carrying pKGB8 or pKGB9 plasmids. Transconjugants were selected on L agar plates with carbenicillin, chloramphenicol and 40 μg ml^-1^ X-gal to allow visualization of *bexR*p-*lacZ* induction immediately after the introduction of plasmids.

### RNA isolation

*P. aeruginosa* strains taken from -80°C stocks were grown on L agar plates at 37°C and single colonies were used to inoculate three independent cultures (biological replicates). After overnight growth, the cultures were diluted 1:100 into fresh L broth and grown to the optical density 0.4- 0.6 at 600 nm (OD_600_). RNA was isolated with RNeasy mini kit (Qiagen) according to the manufacturer’s protocol for bacterial cells from 2 ml of cultures mixed with 4 ml of RNAprotect Bacteria Reagent (Qiagen). Total RNA was digested with DNase (TURBO DNA-free Kit, Ambion) to remove genomic DNA.

### Microarray analysis

Microarray analysis was performed essentially as described before [39]. The raw microarray data have been deposited at the NCBI Gene Expression Omnibus database at accession number GSE95647 (release after publication acceptance). Gene expression data were analysed using Partek Genomic Suite v6.6 (Partek Inc., St. Louis, MO). Raw data were processed using GeneChip Robust Multiarray Averaging (GC RMA): background correction, quantile normalization, log2 transformation and median polish summarization. Analysis of variance (ANOVA) using REML (restricted maximum likelihood) was performed to identify differentially expressed genes. Gene lists were created using a cut-off of *p*-value≤0.05, with a fold change (FC) higher than 2 or lower than -2. Clustering of the genes according to expression pattern changes was performed by K-means clustering [52] using MultiExperiment Viewer v4.9 [53]. The gene expression data for individual replicates were averaged, normalized to zero-mean and unit variance, and subjected to clustering into six clusters, using Pearson correlation as the distance metric and 500 as the maximum number of iterations.

### RT-qPCR analysis

cDNA was synthesized with the TranScriba cDNA synthesis kit (A&A Biotechnology, Poland) using 1.5 μg of total RNA per reaction and random hexamer primers. RT-qPCR reactions were performed in a Roche LightCycler 480 using Hot FIREPol EvaGreen qPCR Mix Plus (Solis Biodyne). The reactions were performed with 0.15- 0.3 μl of cDNA in a total volume of 20 μl. Three technical replicates were used for each gene/primer combination. The primers used to amplify target and reference genes are listed in Table S1. Changes in the gene expression were calculated using the Pfaffl method [54] with normalization to the reference gene *nadB* (*PA0761*). RT-qPCR data represent mean for 3 biological replicates and 3 technical replicates. All RT-qPCR experiments were repeated twice and representative experiments are shown.

### Western blotting

Cultures were grown on L broth to OD_600_ 0.4-0.6. Cells were harvested by centrifugation and stored at -20°C. For each sample a small aliquot was used to estimate the number of c.f.u. ml^-1^ by plating appropriate dilution of the culture on L agar plates. Pellets were resuspended in 10 mM Tris-HCl (pH 8.0), 1 M NaCl, 0.1 mM EDTA, 5% glycerol. For each sample different volume of the buffer was used to compensate the initial differences in c.f.u. ml^-1^. Samples were sonicated and aliquots of cleared extracts corresponding to 10^9^ cells were separated by SDS-PAGE on 12% polyacrylamide gels. Additionally, His_6_-ParB, purified as described [37], was loaded on each gel. Separated proteins were transferred onto nitrocellulose membranes. The blots were subjected to a two-step immunoreaction including application of rabbit polyclonal anti-ParB antibodies followed by incubation with goat anti-rabbit antibodies conjugated with alkaline phosphatase and were developed by addition of NBT/ BCIP mixture. Band intensity was estimated using ImageJ for at least 3 biological replicates.

### Biofilm formation assay

Biofilm formation in static cultures was assayed as described previously with minor modifications [55]. Overnight cultures of *P. aeruginosa* strains in three biological replicates were diluted 1:100 in L broth with or without 0.02% arabinose. 75 μg ml^-1^ chloramphenicol was added to maintain the plasmids. 100 μl aliquots of each diluted culture were transferred into 9 wells of a 96-well flat-bottom polystyrene microplate (Greiner Bio-One). Sterile L broth was used as a negative control. The plate was incubated without shaking at 37°C until OD_600_ reached approximately 0.5. The optical density OD_600_ was measured using a plate reader (BioTek Synergy HT) and bacterial cells, which did not adhere to the surface, were gently removed by aspiration with a pipette. Wells were washed twice with 100 μl of PBS (15 mM KCl, 150 mM NaCl, 10 mM NaPi pH 7.4) and biofilm in each well was stained with 100 μl of 0.1% crystal violet solution. After 15 min incubation at room temperature, crystal violet solution was aspirated and wells were rinsed three times with distilled water and twice with 100 μl of PBS. Subsequently, 100 μl of 96% ethanol was added to each well and left for 10 min to dissolve the crystal violet. Well content was mixed by pipetting and OD_590_ was measured using a plate reader and 96% ethanol as a blank. Biofilm formation is presented as proportion OD_590_/OD_600_.

### β-galactosidase activity assays

β-galactosidase activity was measured as described before [42]. Extracts were prepared from cells grown to OD_600_ 0.4-0.6 in L broth with 75 μg ml^-1^ chloramphenicol, with or without 0.1% arabinose.

## Results

### Impact of *parB* overexpression on gene expression in *P. aeruginosa*

Our previous microarray analysis revealed altered expression of 1166 genes in ParB-deficient cells of *P. aeruginosa* relative to the wild type PAO1161 (WT) cells [39]. To complement this study, here we analysed the transcriptome of ParB -overproducing cells. To control the level of *parB* expression we constructed a pKGB8 expression vector based on the broad-host-range pBBR1MCS-1 [56], which contains the arabinose (Ara) inducible promoter *araBAD*p [49]. Analysis of the growth of PAO1161 (pKGB9 *araBAD*p-*parB*) and PAO1161 (pKGB8) cells in the presence of different concentrations of Ara revealed that induction by Ara ≤0.02% did not affect the growth (Fig 1A). A slight increase in the ParB level (≤2-fold) was observed even in non-induced cells of PAO1161 (pKGB9) relative to the cells containing the empty vector according to Western blot analysis (Fig 1B). Addition of 0.02% Ara to the medium resulted in a 5-fold increase in the ParB amount per cell in mid-log phase cultures of PAO1161 (pKGB9) in comparison to PAO1161 (pKGB8) (Fig 1B). Such ParB excess did not affect the frequency of anucleate cells formation, cell size, colony morphology or swimming and swarming motility (data not shown) but led to altered biofilm formation under specific conditions (Fig 1C, discussed below).

**Fig 1.**
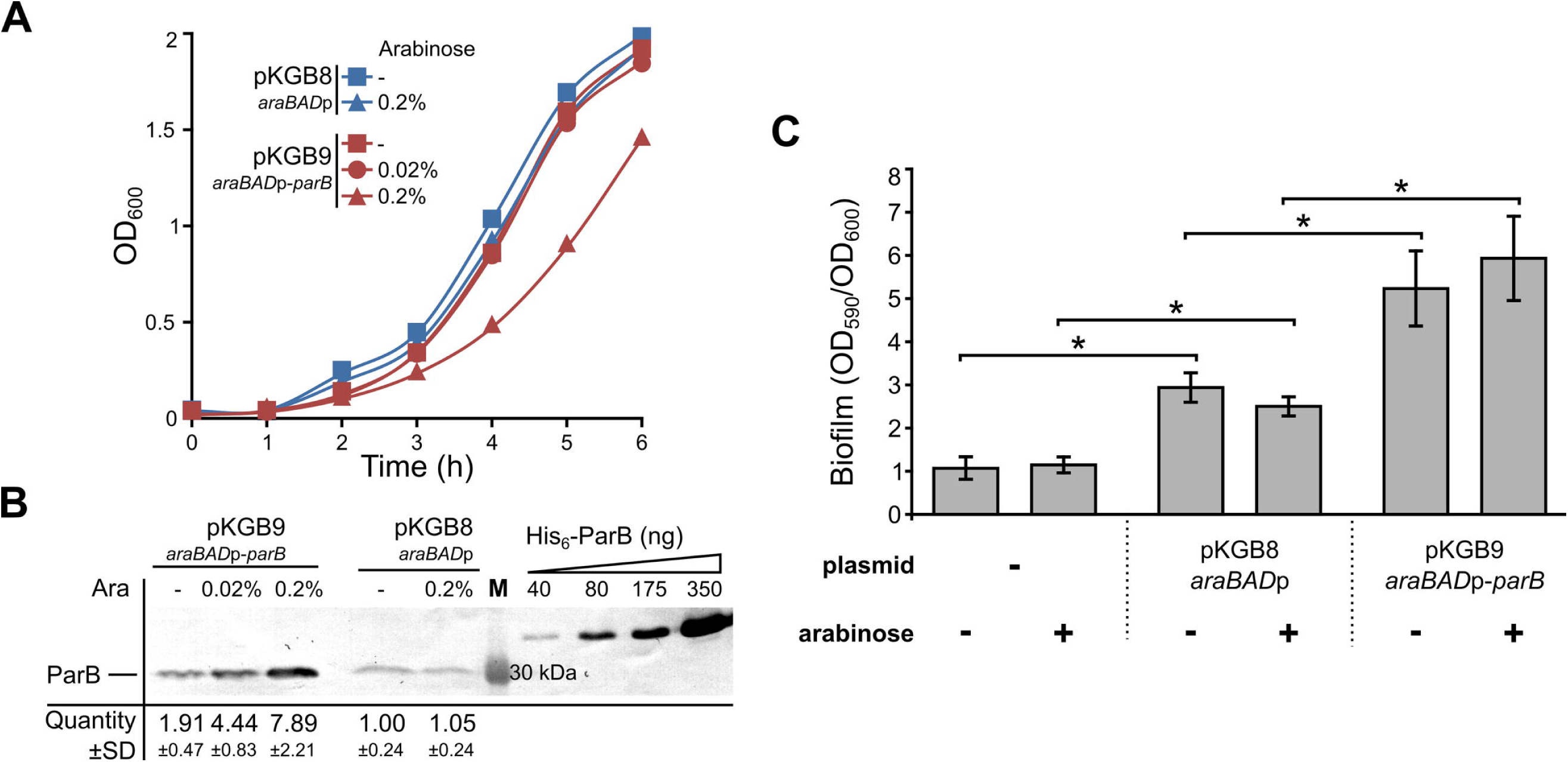
Effects of ParB excess in *P. aeruginosa.* (**A**) Growth of *P. aeruginosa* PAO1161 (pKGB8 *araBAD*p) and PAO1161 (pKGB9 *araBAD*p-*parB*) strains in L broth with different arabinose concentrations. Data represent mean OD_600_. (**B**) Western blot analysis of ParB levels in the tested strains. Each lane contains extract from 10^9^ cells. Blots were subjected to immunodetection using primary anti-ParB antibodies. Representative blot is shown. Signals on the blots were quantified. Data represent mean ParB level ±SD relatively to the control strain PAO1161 (pKGB8). Purified His_6_-ParB was used to generate standard curves. M – molecular weight marker. (**C**) Biofilm formation in the static cultures of PAO1161, PAO1161 (pKGB8) and PAO1161 (pKGB9). Strains were grown without or with 0.02% arabinose until OD_600_ 0.5. Biofilm was stained with crystal violet and assessed by measurement of OD_590_. Data represent mean OD_590_/OD_600_ ratio ±SD from 3 biological replicates. * - *p*-value < 0.05 in two-sided Student’s *t*-test assuming equal variance.

To determine the impact of an increased ParB level on the transcriptome a microarray analysis was performed on RNA isolated from PAO1161 (pKGB9 *araBAD*p-*parB*) cultures grown under selection in L broth with 0.02% Ara (higher ParB overproduction, hereafter referred to as ParB*+++*) or without Ara (mild ParB overproduction, hereafter called ParB*+*) as well as from PAO1161 (pKGB8) cells grown under selection in L broth with 0.02% Ara (empty vector control, hereafter EV) and PAO1161 cells grown in L broth without Ara (wild-type control, hereafter WT). A principal component analysis of the microarray data revealed that the biological replicates for WT and EV form 2 separate groups whereas the biological replicates of ParB*+* and ParB*+++* form together a separate group, distinct from WT and EV samples (data not shown).

Comparative transcriptome analysis of EV *vs* WT revealed 70 genes, four tRNA genes, and four intergenic regions with an altered expression in response to the presence of the vector [fold change (FC) <-2 or >2, *p*-value <0.05] (Fig 2A, Table S2). 175 genes and 1 intergenic region displayed statistically significant difference in expression between ParB*+++* and EV cells using the same criteria. 155 out of 175 genes displayed an altered expression only in the response to the ParB abundance since they were not identified in the EV *vs* WT analysis. The expression of the remaining 20 genes and one intergenic region was affected by the presence of vector as well as ParB overproduction in ParB+++ cells (Fig 2B, Table S2). The comparative analysis of ParB*+* and EV transcriptomes revealed altered expression of 157 genes. There was a major overlap between the transcriptomic changes in ParB+ and ParB+++ cells as 122 genes showed similar responses in the ParB+ and ParB+++ cells (Fig 2B, Table S2). This is not surprising as the transcription of *parB* (*PA5562*) itself was increased 4- and 12- fold in ParB*+* and ParB*+++* cells, respectively, in comparison to the EV cells.

**Fig. 2.**
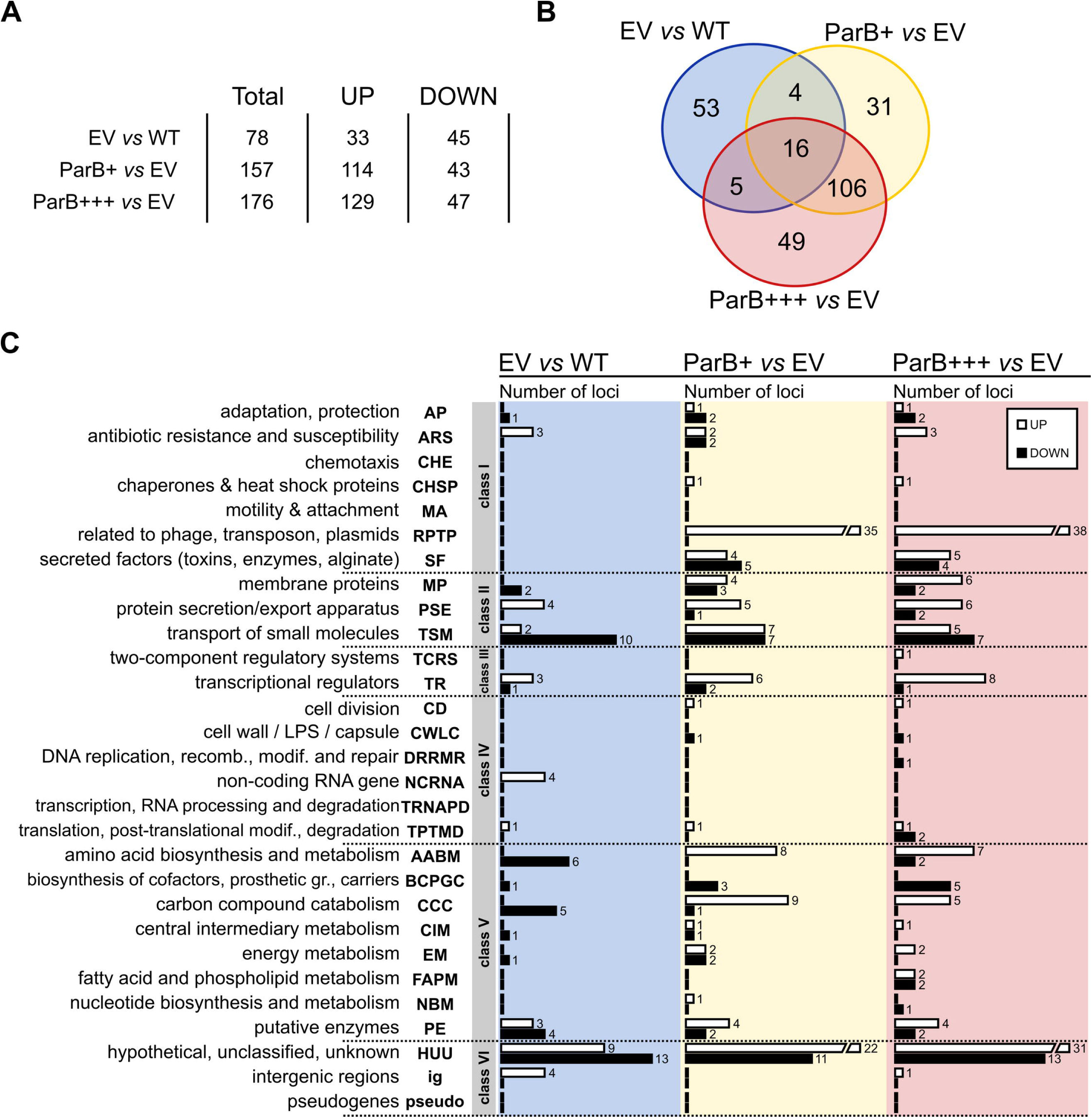
Transcriptome changes in response to ParB overproduction. (**A**) Statistics of loci with significant expression change (FC<-2 or >2, *p*-value <0.05). RNA was isolated from PAO1161 cultures grown in L broth without Ara (WT), PAO1161 (pKGB8 *araBAD*p) cultures grown under selection in L broth with 0.02% Ara (empty vector control, EV), PAO1161 (pKGB9 *araBAD*p-*parB*) cultures grown under selection in L broth without Ara (mild ParB excess, ParB+) or with 0.02% Ara (higher ParB excess, ParB*+++*). (**B**) Venn diagram for sets of loci with significant expression change between EV *vs* WT, ParB+ *vs* EV and ParB+++ *vs* EV. (**C**) Classification of loci with altered expression according to PseudoCAP categories [57]. When a gene was assigned to multiple categories, one category was arbitrarily selected (Table S2). The PseudoCAP categories were grouped into six classes as marked. White and black bars correspond to the numbers of respectively, upregulated and downregulated genes in a particular category.

Interestingly, there was also a set of 35 genes with expression changed only in ParB+ cells relative to EV cells but not in ParB+++. Closer inspection of the expression changes of these genes revealed three subgroups (Fig S1A, Table S2): i/ 15 genes were also altered in ParB+++ transcriptome but with a slightly lower fold change (FC>1.6 or FC<-1.6*, p*-value <0.05) and were cut off from the common pool of ParB -affected genes using the criterion of FC, ii/ 6 genes despite their high FC were eliminated in ParB+++ vs EV analysis because of slightly higher variation of their expression in biological replicates (0.05< *p*-value <0.1), iii/ remaining 14 genes seemed to be not altered in ParB+++ transcriptome. Whereas ParB+++ and EV strains were grown under the same conditions (chloramphenicol and 0.02% arabinose), ParB+ cells were not exposed to arabinose. To estimate the impact of inducer on gene expression PAO1161 (pKGB8) was grown in medium with or without 0.02% Ara and RT-qPCR analysis of eight loci from the group of 35 genes in question, representing all three subgroups, was performed. The expression of only one gene, *PA2113* (from the bi-cistronic operon *PA2113-2114*) was altered in response to Ara presence in the medium (Fig S1B). *PA2113-2114* belong to the third subgroup in the expression pattern analysis (see above) but also to a distinct set of 4 genes common for EV *vs* WT and ParB+ *vs* EV lists (Fig 2B, Table S2), hence it is likely that these four genes appeared in our analysis due to the influence of Ara on their expression. Presence of Ara had no influence on the expression of 7 out of 8 genes tested, which represent 14 genes due to the operon structures (Fig S1B), confirming that majority of genes with the expression altered in ParB+ cells relative to EV cells was identified due to their response to ParB excess.

The genes with altered expression were assigned to PseudoCAP categories [57] divided arbitrarily into six classes as described previously [39]. The comparison of the EV and WT transcriptomes revealed that the majority of affected genes belong to three groups: Class II comprising genes encoding membrane proteins, proteins involved in transport of small molecules, and protein secretion systems (18 genes), Class V comprising genes involved in metabolic pathways (21 genes), and Class VI with 22 HUU genes (hypothetical, unclassified, unknown) (Fig. 2C, Table S2). Similarly, most of the genes with altered expression in ParB overproducing cells (ParB+++ and ParB+) relative to EV cells were assigned to the same three classes (II, V and VI). However, almost one third of the genes with expression altered in response to ParB overproduction fall into Class I, mainly to PseudoCAP categories RPTP (related to phage, transposon, plasmids) and SF (secreted factors: toxins, enzymes, alginate). These data suggest that an increased ParB level leads to the activation of stress response.

### Different expression patterns of genes affected by ParB excess

To systematize the impact of the vector as well as the increased level of ParB on the PAO1161 transcriptome a K-means clustering analysis was performed [52]. The two lists of altered genes (ParB+ vs EV and ParB+++ vs EV) were combined yielding 211 unique loci (Table S2). Expression data for replicates for each of these loci in the four strains/ growth conditions (WT, EV, ParB+, ParB+++) were averaged, normalized to zero mean and unit variance and grouped into six clusters based on the similarity of their expression patterns (Fig 3A and B).

**Fig 3.**
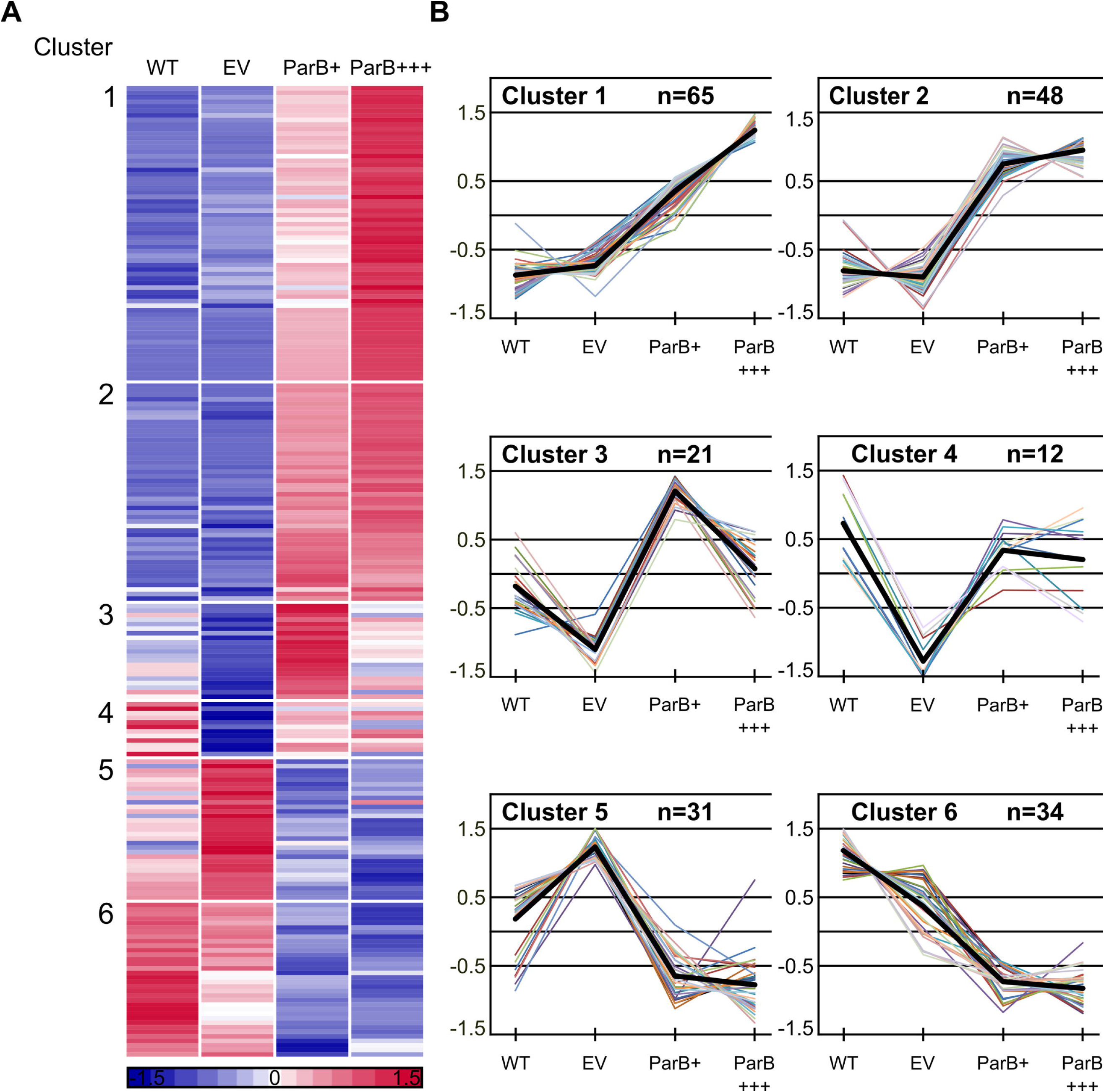
K-means clustering of microarray data. Expression data for replicates for each of 211 loci displaying altered expression in ParB overproducing cells were averaged, normalized to zero mean and unit variance and grouped into six clusters. (**A**) Expression profiles of individual genes grouped according to the results of K-means clustering. Each horizontal line represents one gene. Red and blue denote that the expression is respectively, above or below the mean expression of a gene across the data set. (**B**) Expression profiles for genes in each cluster. Y-axis represents the difference between expression of a particular gene in tested conditions and the mean expression of this gene in all 4 conditions presented as the number of standard deviations that a particular data point differs from the mean. Thick black lines represent the cluster centres. The genes from each cluster and their expression levels in different cells are listed in Table S3.

Clusters 1 to 4 contain genes that are upregulated in ParB*+* and/or ParB+++ cells relative to EV cells and clusters 5 and 6 the downregulated ones. A detailed description of the genes in each cluster as well as their mean expression levels in the four conditions are given in Table S3.

To validate the expression patterns obtained from the microarray analysis the transcript level for selected genes from each cluster in various samples was analysed using RT-qPCR. Importantly, for all tested 12 genes their relative levels of expression detected by RT-qPCR correlated nearly perfectly with the microarray data (Table 3).

**Table 3.** Validation of the microarray data by RT-qPCR.

| Cluster | Gene ID | WT |  | EV | ParB<sup>+</sup> |  | ParB<sup>+++</sup> |  |
| --- | --- | --- | --- | --- | --- | --- | --- | --- |
|  |  | MA | qPCR | qPCR | MA | qPCR | MA | qPCR |
| 1 | *PA0612* | 0.66 | 0.76 ± 0.22 | 1 ± 0.06 | 6.54 | 5.62 ± 0.97 | 10.57 | 9.69 ± 3.3 |
| 1 | *PA0985* | 0.83 | 0.87 ± 0.16 | 1 ± 0.32 | 3.57 | 5.7 ± 1.67 | 5.02 | 7.1 ± 2.14 |
| 1 | *PA4635* | 0.60 | 0.48 ± 0.08 | 1 ± 0.12 | 2.95 | 1.42 ± 0.47 | 4.63 | 2.39 ± 1.12 |
| 1 | *PA5562* | 0.93 | 0.94 ± 0.21 | 1 ± 0.08 | 4.31 | 2.49 ± 0.43 | 12.06 | 17.3 ± 3.21 |
| 2 | *PA2432* | 0.62 | 0.57 ± 0.18 | 1 ± 0.29 | 8.36 | 5.13 ± 1.13 | 7.90 | 6.74 ± 0.1 |
| 2 | *PA3661* | 1.16 | 1.15 ± 0.45 | 1 ± 0.18 | 17.50 | 5.78 ± 2.23 | 23.34 | 8.39 ± 4.41 |
| 2 | *PA5471* | 0.24 | 0.28 ± 0.08 | 1 ± 0.25 | 2.24 | 2.01 ± 0.66 | 2.50 | 2.34 ± 0.34 |
| 3 | *PA2008* | 2.44 | 3.03 ± 1.06 | 1 ± 0.27 | 6.26 | 7.91 ± 1.36 | 3.81 | 3.91 ± 0.09 |
| 4 | *PA4659* | 1.73 | 0.97 ± 0.13 | 1 ± 0.13 | 1.83 | 2.23 ± 0.22 | 2.13 | 2.06 ± 0.1 |
| 5 | *PA0996* | 0.75 | 0.57 ± 0.01 | 1 ± 0.15 | 0.58 | 0.63 ± 0.22 | 0.31 | 0.33 ± 0.04 |
| 5 | *PA1003* | 0.87 | 0.73 ± 0.13 | 1 ± 0.24 | 0.72 | 0.58 ± 0.05 | 0.45 | 0.44 ± 0.04 |
| 6 | *PA0011* | 0.96 | 1.01 ± 0.2 | 1 ± 0.24 | 0.37 | 0.41 ± 0.03 | 0.20 | 0.24 ± 0.00 |

Relative transcript level of selected genes in replicates of *P. aeruginosa* cultures assessed by microarrays and RT-qPCR. WT - PAO1161 grown in L broth, EV - PAO1161 (pKGB8) grown in L broth with 0.02% Ara,
ParB<sup>+</sup> - PAO1161 (pKGB9) grown in L broth and ParB<sup>+++</sup> - PAO1161 (pKGB9) grown in L broth with 0.02% Ara. Microarray (MA) data represent mean expression and RT-qPCR (qPCR) data represent mean ± SD from three biological replicates.

### Genes upregulated in response to *parB* overexpression

Clusters 1 (64 genes and one intergenic region), 2 (48 genes), 3 (21 genes) and 4 (12 genes) contain loci upregulated in response to ParB overproduction. For eight of these loci an increased transcript level is also observed in EV cells relative to WT cells (Table S3, Table 4). Among these loci are *PA5471* gene encoding ArmZ, an inhibitor of MexZ which negatively regulates the *mexXY* operon encoding the multidrug efflux pump MexXY [58,59] and *pyeR* (*PA4354*) encoding a transcriptional regulator of biofilm formation [60]. Additionally, Cluster 1 contains the *arr* (*PA2818*) gene predicted to encode a phosphodiesterase whose substrate is cyclic di-guanosine monophosphate (c-di-GMP), a bacterial secondary messenger that regulates cell surface adhesiveness, virulence and biofilm formation [61].

**Table 4.**
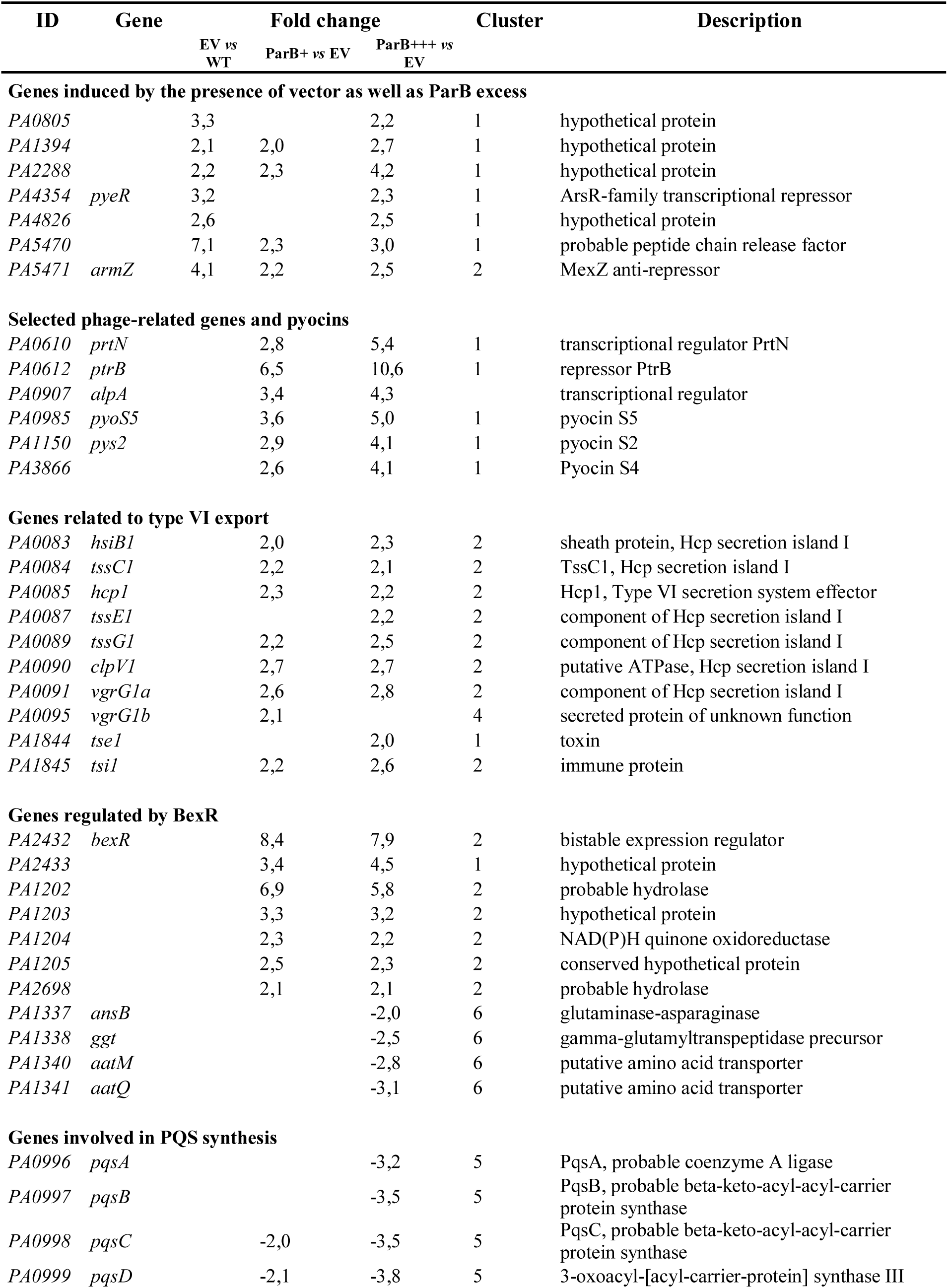
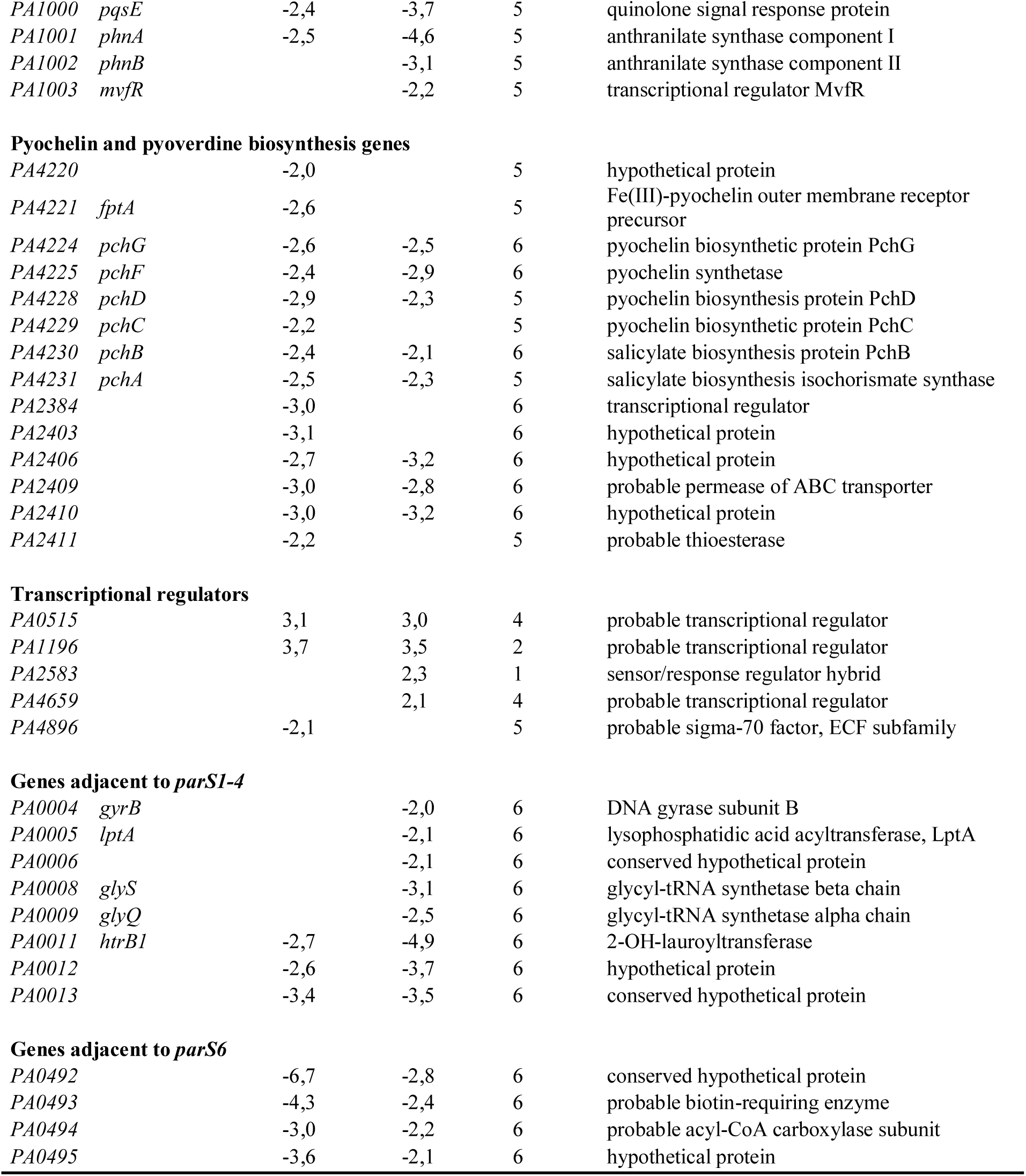
Selected genes whose expression was identified as ParB regulated.

The biofilm formation was checked in the cultures of PAO1161 (pKGB9 *araBAD*p-*parB*) in comparison to PAO1161 (pKGB8) during growth under static conditions. No difference was observed between the strains in the overnight cultures (data not shown). Notably, a significant increase in biofilm formation was observed in dividing cultures of PAO1161 (pKGB9 *araBAD*p-*parB*) relative to PAO1161 (pKGB8), which correlates with the altered expression of genes involved in regulation of biofilm formation and identified in our study. No effect of arabinose presence on biofilm formation was detected (Fig 1C). Transcriptomic analysis also implicated that ParB excess could alter the response to antibiotic presence. The MICs for six antibiotics from β-lactam, aminoglycoside and fluorochinolone groups (piperacillin, ticarcillin, imipenem, gentamicin, tobramycin and ciprofloxacin) were tested but no significant differences between PAO1161 (pKGB9) and control PAO1161 (pKGB8) strains grown with and without arabinose were detected (data not shown).

A remarkable feature of clusters 1 and 2, comprising the majority of ParB -upregulated genes (Table S3), is a high proportion of phage-related genes (*PA0610, PA0612-0641, PA0643-0648*, *PA0717*, *PA0718*, *PA0907*, *PA0909-0911*) as well as pyocin-encoding genes (*PA0985*, *PA1150*, *PA3866*). Induction of bacteriophage genes and pyocin production in *P. aeruginosa* have been shown to be a hallmark of the SOS response [62], suggesting that *parB* overexpression induces a DNA damage signal. In this bacterium SOS response is coordinated not only by LexA (*PA3007*), but also by two structurally related repressors, PrtR (*PA0611*) and AlpR (*PA0906*) [62,63]. Significantly, expression of none of these genes is affected by ParB excess. PrtR is an inhibitor of *prtN* (*PA0610*), a transcriptional activator of pyocin synthesis genes [64]. *prtN* is induced 5.4-fold in ParB+++ cells suggesting a relief of the PrtR-mediated repression in these cells. Similarly, genes controlled by AlpR (*PA0907*, *PA0909-PA0911*) are induced in response to ParB overproduction. Further studies are required to determine the effect on the expression of the PrtR- and AlpR- regulated genes apparently without a change in the amounts of transcript level of these two repressors.

The ParB excess also induces genes encoding components of the Hcp secretion island HSI-1 [65] (*PA0083-0085, PA0087, PA0089-0091*) and the toxin/immunity proteins Tse1 (*PA1844)* and Tsi1 (*PA1845*). The HSI-1 together with two other secretion islands participate in *P. aeruginosa* virulence, inter- and intraspecies antagonism, biofilm formation, and stress sensing [66].

Overexpression of *parB* also results in an 8-fold increase in the expression of the transcriptional regulatory gene *PA2432* encoding a bistable response regulator BexR [67]. Six genes, *PA1202, PA1203, PA1204, PA1205, PA2433* and *PA2698*, previously identified as upregulated in response to BexR overproduction [67], are also upregulated in ParB-overproducing strains. Similarly, five genes downregulated in response to BexR overproduction, *PA0998, PA1337, PA1338, PA1340* and *PA1341*, are also downregulated in the analysed ParB -overproducing cells (Table 4).

The expression of *bexR* gene is known to be bistable, meaning that this gene switches between OFF and ON states in cells of a genetically identical bacterial population [67]. The microarray analysis was performed on three independent biological isolates of PAO1161 (pKGB9 *araBAD*p-*parB*) and all demonstrated a strong induction of *bexR* (*PA2432*) expression (*p*-value 3E-06) strongly suggesting that ParB excess modulates the BexR regulon through induction of *bexR* expression. To monitor the effect of ParB excess on promoter of *bexR* the strain PAO1161::*bexR*p-*lacZ* was constructed with the promoter-reporter cassette inserted in a non-coding region of the genome. A single colony (white on L agar with X-gal) was inoculated and the cells were used as the recipients in conjugation with either S17-1 (pKGB8) or S17-1 (pKGB9 *araBAD*p-*parB*). Conjugants were plated on selective medium containing X-gal to visualize the expression of *bexR*p-*lacZ* transcriptional fusion. Conjugants PAO1161::*bexR*p-*lacZ* (pKGB8) formed typical white/transparent colonies with a very low frequency of blue ones (less than 0.1%) whereas majority of PAO1161::*bexR*p-*lacZ* (pKGB9) conjugants formed blue colonies (Fig 4), confirming that ParB induces *bexR*p expression.

**Fig 4.**
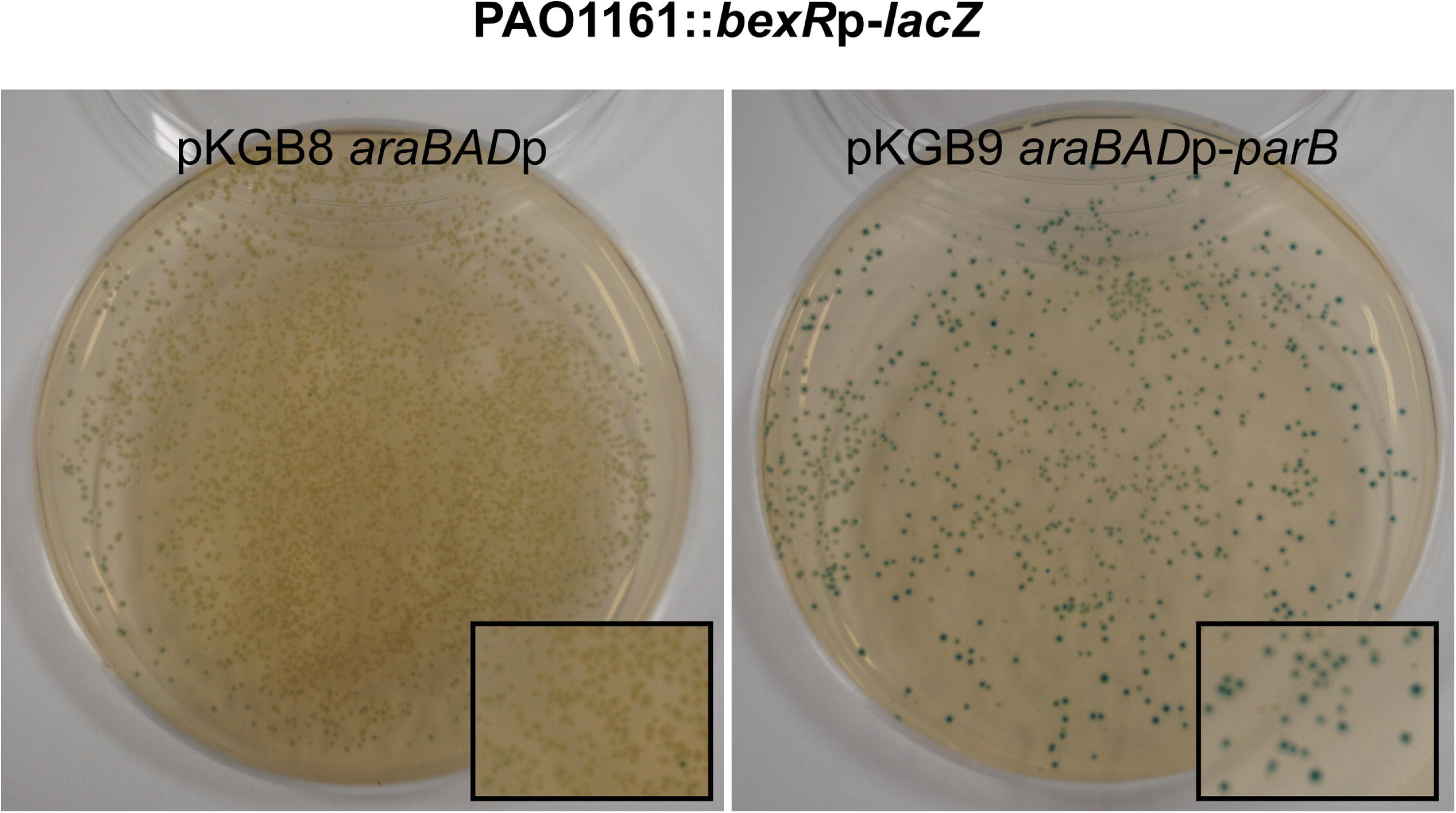
ParB excess induces the expression of chromosomal *bexR*p-*lacZ* transcriptional fusion. PAO1161::*bexR*p-*lacZ* strain contains *bexR*p-*lacZ* transcriptional fusion inserted in the intergenic region of PAO1161 genome. White colony from L agar with X-gal was inoculated and used as a recipient in conjugation with either S17-1 (pKGB8) or S17-1 (pKGB9 *araBAD*p-*parB*) donor cells. Conjugants were grown on selective L agar plates supplemented with X-gal. The photographs show representative plates of PAO1161::*bexR*p-*lacZ* with both plasmids.

Clusters 3 and 4 group 33 genes with mRNA level reduced in response to plasmid/ chloramphenicol/ arabinose (EV) but then increased by ParB excess. Twelve of the 33 genes in this group belong to the PseudoCAP category carbon compounds catabolism (CCC) and eight genes to the amino acid biosynthesis and metabolism (AABM) category (Table S3). The altered expression of genes from these categories suggests that ParB overproduction may interfere with primary metabolism as an adaptive measure. Interestingly, three out of four putative arabinose-dependent genes have also been classified into cluster 4. Additionally, cluster 4 contains two putative transcriptional regulators: *PA0515*, which is a part of the *nir* operon (denitrification operon), and *PA4659* with unknown function.

### Genes downregulated in response to *parB* overexpression

Clusters 5 (31 genes) and 6 (34 genes) contain genes downregulated in response to ParB overproduction but differ from each other by a moderate induction in EV cells seen in Cluster 5. This cluster contains all genes from two operons, *pqsABCDE* (*PA0996-1000*) and *phnAB* (*PA1001* and *PA1002*), involved in the production of *Pseudomonas* quinolone quorum sensing signal (PQS) [68,69]. Interestingly, the same cluster contains *PA1003* (*mvfR/pqsR*) which encodes a positive regulator of the both operons [70,71], suggesting that the downregulation of the *pqs* and *phn* operons is a direct result of lower *mvfR* expression. Interestingly, MvfR also acts as a negative regulator of type VI secretion system HSI-1 [66], components of which are upregulated in response to ParB excess (Table 4 and see above).

Five genes (*PA4220*, *PA4221*, *PA4228*, *PA4229*, *PA4231*) from Cluster 5 and three genes (*PA4224*, *PA4225*, *PA4230*) from Cluster 6 are involved in the biosynthesis and transport of a siderophore, pyochelin. Pyochelin synthesis in *P. aeruginosa* cells is positively regulated by the transcriptional regulator PA2384 [72], which is downregulated in ParB+ cells. Interestingly, biosynthesis of another siderophore, pyoverdine, also seems to be negatively affected by ParB excess as the genes *PA2403*, *PA2406*, *PA2409* and *PA2410* (Cluster 6) from the pyoverdine biosynthesis operon are significantly downregulated in ParB-overproducing cells (Table 4).

### Analysis of ParB-related transcriptional silencing around *parS* sites

Analysis of the microarray data revealed that ParB overproduction reduces the expression of a number of genes in close proximity of *parS1-4* (Table 4). Closer inspection of the expression changes in this region revealed that genes *PA0003-PA0015*, with the exception of *PA0007*, *tag* (*PA0010*) and *PA0014*, show a significant (30%-80%, *p*-value <0.05), ParB dose-dependent downregulation (Fig 5A). RT-qPCR analysis revealed that in fact all genes from *PA0004* to *PA0014* are subject to significant downregulation (Fig 5B), suggesting that ParB bound to *parS1*-*4* negatively influences the expression of adjacent genes. To verify this hypothesis we analysed the expression of these genes in the *parB*_null_ mutant, which does not produce ParB [18], and in the *parS*_null_ mutant in which the *parS1-4* sequences had been mutated, so the ParB binding to these sequences was impaired [38]. In the both mutants only genes from *PA0010* to *PA0015* displayed significantly increased expression (Fig 5C) indicating that at its native level ParB acts as a major negative regulator of genes adjacent to intergenic *parS3* and *parS4* but not for the genes adjacent to intragenic *parS1* and *parS2*. Regulation of the expression of genes in *parS1-parS4* region seems to be a direct consequence of ParB binding to *parS1*-*parS4* as ParB overproduction in *parS*_null_ cells did not lower the expression of *dnaA*–*trkA* (*PA0001*-*PA0016*) genes relatively to the control *parS*_null_ cells carrying empty vector (Fig 5D, also see below).

**Fig 5.**
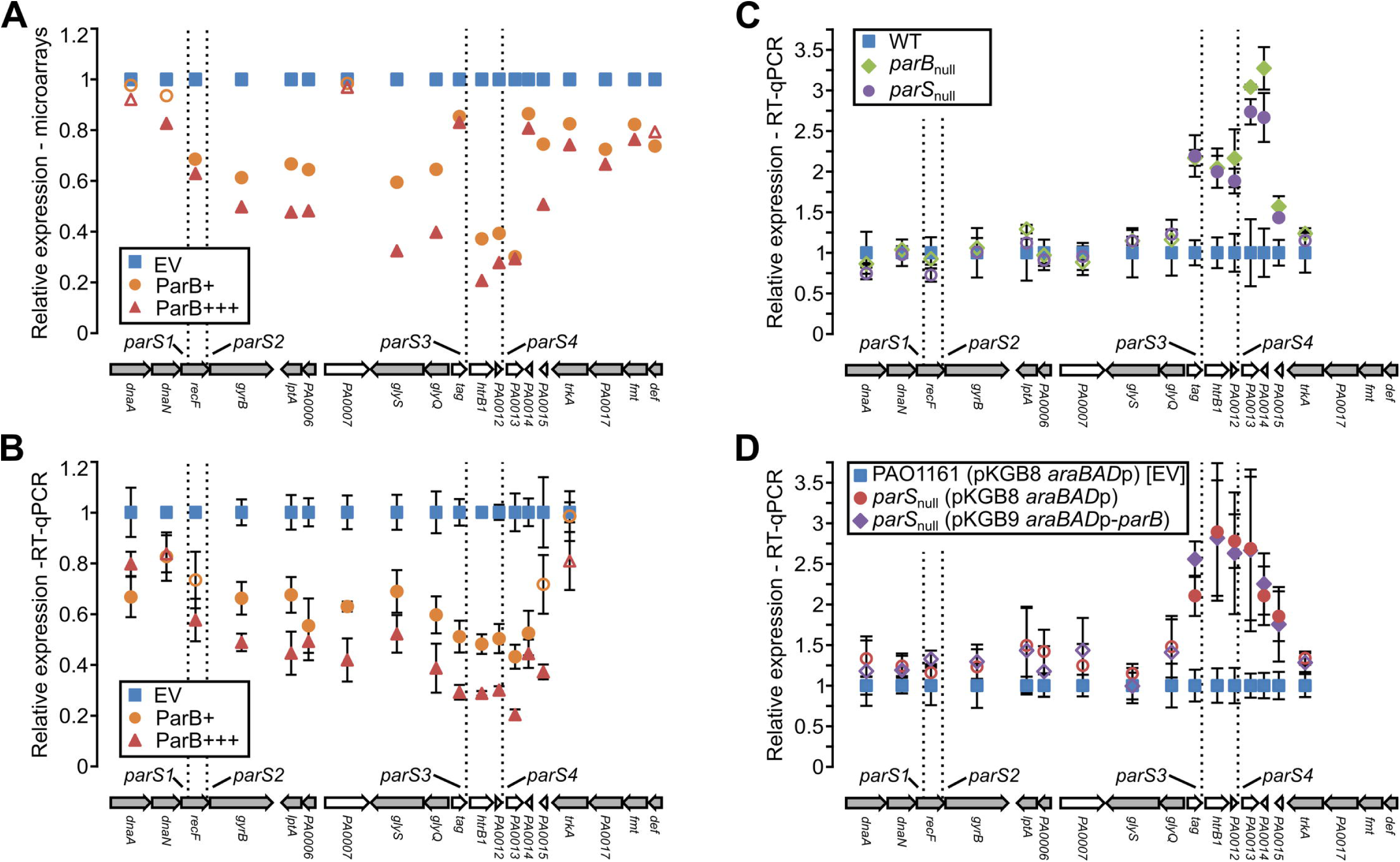
Influence of ParB on gene expression in the *parS1-4* region. (**A**) Mean level of expression of *dnaA*-*def* (*PA0001*-*PA0019*) genes in ParB+ and ParB+++ cells relative to EV as revealed by microarray analysis. Filled markers indicate statistically different expression relative to EV (*p*-value < 0.05 in ANOVA test). Arrangement of the genes in the chromosome is shown below. Operons (according to the DOOR 2.0 database [73]) are marked in grey. (**B**) Expression of *PA0001*-*PA0016* (*dnaA*–*trkA*) genes in ParB+ and ParB+++ cells relative to EV cells. (**C**) RT-qPCR analysis of expression of *dnaA*–*trkA* genes in *parB*_null_ and *parS*_null_ strains relative to WT cells. Cells were grown in L broth. (**D**) RT-qPCR analysis of expression of *dnaA*–*trkA* genes in *parS*_null_ (pKGB9 *araBADp*-*parB*) and *parS*_null_ (pKGB8 *araBAD*p) relative to EV [PAO1161 (pKGB8 *araBAD*p)]. Cells were grown in L broth supplemented with chloramphenicol and 0.02% arabinose. RT-qPCR data represent mean ±SD from three biological replicates. Filled symbols indicate significantly different expression (*p*-value <0.05 in two-sided Student’s *t*-test assuming equal variance) relative to the control cells labelled as blue squares. The differences in expression of genes in *parS*_null_ (pKGB9) strain relative to *parS*_null_ (pKGB8) strain are not statistically significant.

A similar inspection of the expression changes in proximity (±10 adjacent genes) of the remaining six *parS* sequences [38] revealed that overproduction of ParB lowers the expression of genes adjacent to *parS6* (Fig 6A, *PA0492-PA0496*) but not to the other *parS* sequences (data not shown). These results were confirmed by RT-qPCR (data not shown). However, the expression of *PA0492*, *PA0493* and *PA0494* as assayed by RT-qPCR was not significantly changed in *parB*_null_ or *parS*_null_ cells in comparison with the WT strain (Fig 6B), indicating that at its native abundance ParB is not a major effector of genes in this region.

**Fig 6.**
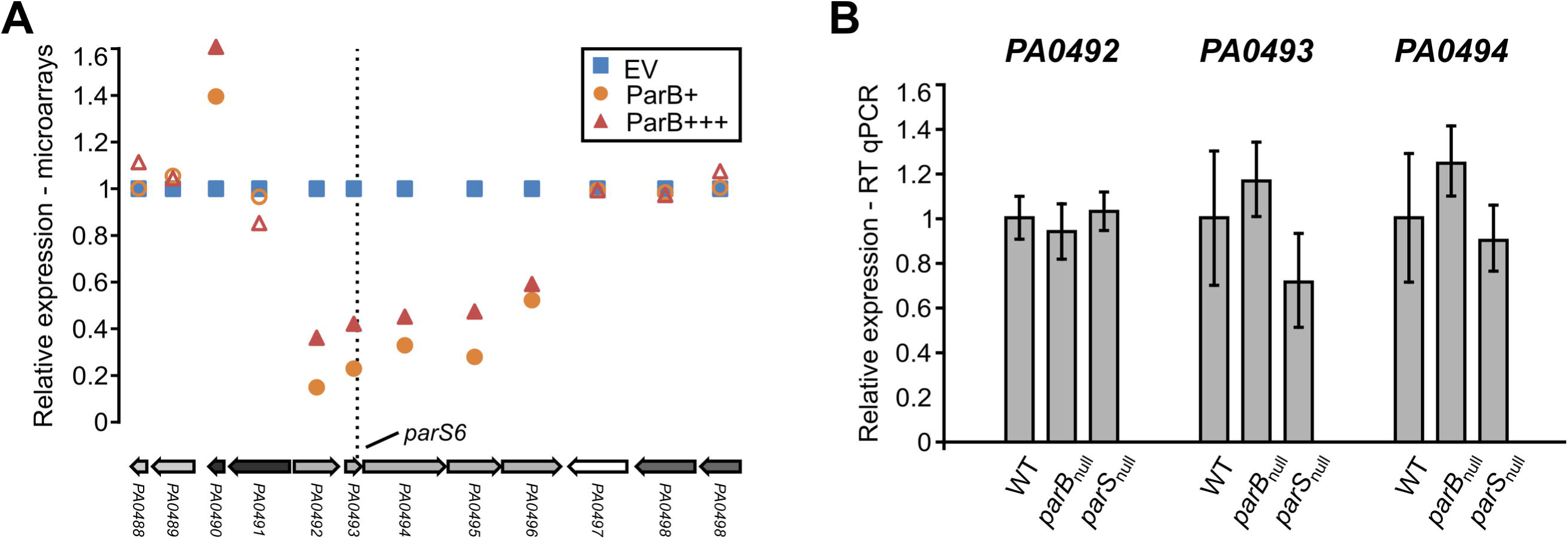
Influence of ParB level on expression of genes adjacent to *parS6*. (**A**) Mean level of expression of *PA0488*–*PA0498* genes in ParB+ and ParB+++ cells relative to EV cells (microarray data). Filled markers indicate statistically different expression relative to EV (*p*-value < 0.05 in ANOVA test). Operons (according to the DOOR 2.0 database [73]) are labelled with different shades of grey. (**B**) RT-qPCR analysis of expression of *PA0492*, *PA0493* and *PA0494* genes in *parB*_null_ and *parS*_null_ strains relative to WT cells. Data represent mean ±SD from three biological replicates. The differences between strains / conditions are not statistically significant (*p*-value > 0.05 in two-sided Student’s *t*-test assuming equal variance).

### Regulation of *PA0011* and *PA0013* promoters by ParB interactions with *parS3* and *parS4* sequences

To further confirm that ParB binding to *parS3* and *parS4* affects the expression of adjacent genes the intergenic regions preceding *PA0011* and *PA0013* and their respective variants carrying mutated *parS3* and *parS4* (Fig 7A) [38] were cloned upstream of a promoter-less *lacZ* cassette in pPJB132, a derivative of pCM132 [50]. To produce an excess of ParB from the chromosome strain PAO1161::*araBAD*p-*flag-parB* was constructed with the expression cassette inserted in a non-coding region of the genome (Table 1). Growth of the PAO1161::*araBAD*p-*flag-parB* cells in a medium containing 0.1% arabinose results in the overproduction of Flag-ParB to a level that is not toxic for the cells (data not shown). All four pPJB132 derivatives were introduced into PAO1161::*araBAD*p-*flag-parB* as well as into control strain PAO1161::*araBAD*p with the empty expression cassette inserted in the same genomic position.

**Fig 7.**
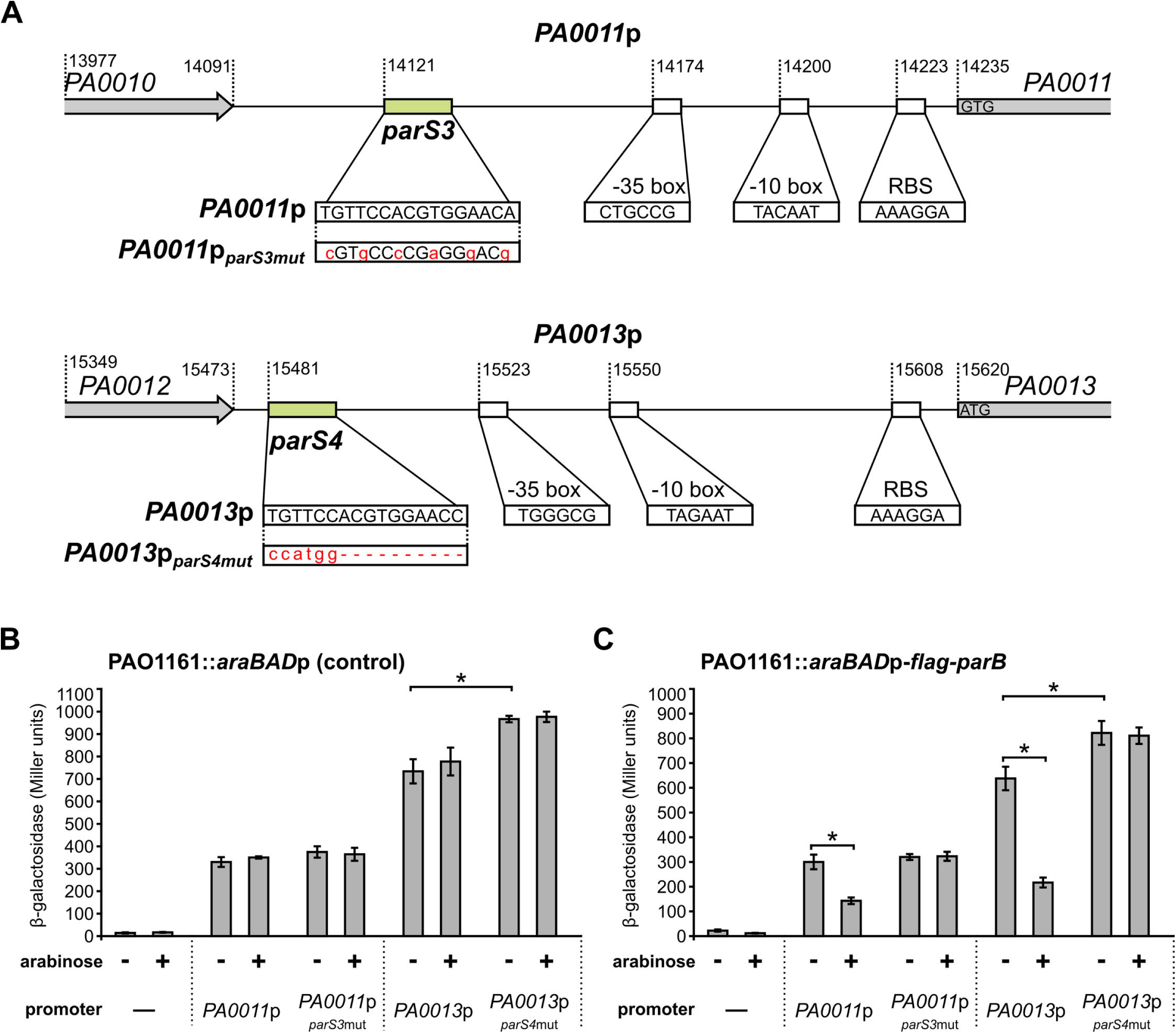
Influence of ParB on the activities of *PA0011* and *PA0013* promoters. (**A**) DNA sequences preceding *PA0011* (*PA0011*p) and *PA0013* (*PA0013*p) and their mutated versions, *PA0011*p*_parS3_*_mut_ and *PA0013*p*_parS4_*_mut_, cloned upstream of promoter-less *lacZ* cassette in pPJB132 are presented. Putative promoters’ motifs, RBS sequences and start codons are indicated. Promoter -35 and -10 boxes were predicted using BPROM [74]. β-galactosidase activity was measured in extracts from PAO1161::*araBAD*p (control strain) (**B**) and PAO1161::*araBAD*p-*flag*-*parB* (ParB-overproducing strain) (**C**) cells carrying pPJB132 derivatives as indicated. Data represent mean activity from at least three cultures ±SD. * - *p*-value < 0.05 in two-sided Student’s *t*-test assuming equal variance.

Analysis of the β-galactosidase activity in the transformants of the control strain revealed no effect of 0.1% arabinose on the promoters tested (Fig 7B). The nucleotide substitutions in *parS3* modifications (Fig 7A) did not affect the *PA0011*p activity (Fig 7B). Surprisingly, replacing *parS4* by a NcoI restriction site (deletion of 10 bp) resulted in a minor increase of the promoter strength in the *PA0013*p*_parS4_*_mut_*-lacZ* fusion (Fig 7B). DNA sequence analysis of promoters’ regions does not give a clear explanation of such effect (Fig 7A). The β-galactosidase activities detected in transformants of PAO1161::*araBAD*p-*flag*-*parB* were similar to those in the corresponding control strain and when the strains were grown without arabinose. However, upon induction of *flag*-*parB* expression with 0.1% arabinose a reduction of β-galactosidase activity was observed in PAO1161::*araBAD*p-*flag*-*parB* transformants carrying plasmids with the *PA0011*p-*lacZ* and *PA0013*p-*lacZ* transcriptional fusions but not plasmids with the *PA0011*p*_parS3_*_mut_-*lacZ* or *PA0013*p*_parS4_*_mut_-*lacZ* transcriptional fusions (Fig 7C). These data confirmed that ParB negatively regulates expression of these two promoters *in vivo* through interactions with *parS* sites.

## Discussion

In *P. aeruginosa*, partitioning protein ParB plays a non-essential but important role in segregation of chromosomes. Its binding to at least one of the four *parS* sequences in the *parS1-4* cluster, closest to *oriC,* is necessary and sufficient for accurate segregation [33,38]. The role of the remaining six *parS* sequences (four in the *ori* domain and two, *parS7* and *parS8*, close to the terminus domain [38]) is not fully understood. Ten similarly distributed *parS* sequences in *B. subtilis* have been shown to participate in SMC recruitment, condensation and juxtapositioning of chromosome arms [75]. Several reports have indicated that partitioning proteins may also modulate transcription of certain genes [34,36] but *P. aeruginosa* seems to be unique, as inactivation of *parA* or *parB* leads to large-scale changes of its transcriptome [39]. A lack or an excess of ParB protein is manifested by similar phenotypic changes, including erratic chromosome segregation, increased rate of formation of anucleate cells, cell elongation, disturbed division and defects in swarming and swimming [17,18,37]. To find out how such multiple effects arise, a microarray-based transcriptomic analysis was performed of PAO1161 derivatives in which ParB level was increased either 2- or 5- fold in comparison with the strain carrying empty vector. Since after binding to a specific *parS* sequence ParB has the ability to spread on the adjacent DNA, we mainly focused on expression of genes in the vicinity of *parS* sequences.

At tested ParB concentrations its binding to *parS* sites has no pronounced effect on expression of genes around them with the exception of genes located in proximity of *parS1-4* and *parS6*. Microarray and RT-qPCR data show convincingly that all the genes adjacent to *parS1-4* (*PA0003* to *PA0015*) are downregulated in the presence of ParB excess (Fig 3A and B). The interference of ParB with the expression of a vital replication operon (*dnaA-dnaN-recF-gyrB*), likely caused by ParB binding to *parS1*/*parS2* sites within the *recF* gene, may underlie the defects in cell division caused by a ParB excess larger than studied here. In contrast, in cells lacking a functional ParB protein or carrying mutated *parS1-4* sequences unable to bind ParB, RT-qPCR analysis revealed only upregulation of *PA0010*-*PA0015* genes, located in proximity of the intergenic *parS3* and *parS4.* Linking the regions preceding *PA0011* and *PA0013* to the promoter-less *lacZ* cassette confirmed the presence of functional promoters in the cloned fragments. Both promoters were repressed by ParB excess provided functional *parS3* and *parS4* were present, which confirmed that ParB binding directly affects transcription in this region.

The product of *PA0011*, the 2-OH-lauroyltransferase HtrB1 is involved in lipid A biosynthesis important for integrity of the outer membrane and cell envelope [76–78]. *htrB1* expression is induced by sub-inhibitory concentrations of different antibiotics [78] or growth at low temperature [77]. Inactivation of *htrB1* leads to pleiotropic effects, manifested by increased susceptibility to surfactants and antibiotics, impaired swarming motility, growth defects and induction of type III secretion system. Some of these effects are also observed under ParB excess. Conversely, expression of another HtrB homolog (*PA3242*) is significantly reduced in *parB*_null_ mutant [39]. Since inactivation of either of the *htrB* genes in *P. aeruginosa* PAO1 leads to motility defects [77,79], this could explain the impaired motility seen both when ParB is missing and when it is present in excess. Notably, also other genes adjacent to *parS3* and *parS4* encode proteins with putative roles in response to cellular stress. *tag* (*PA0010*) encodes a putative 3-methyladenine glycosidase I [80]. *PA0012* encodes a protein with a predicted PilZ domain, which could bind cyclic di-GMP, an important signalling molecule in *P. aeruginosa* [81,82]. Further studies are needed to characterize the biological functions of all the genes adjacent to *parS3* and *parS4* and the role of ParB in their regulation.

Apart from the genes in proximity to *parS3/*-*parS4*, also those close to *parS6* showed a reduced expression in response to ParB excess, however, their expression was not altered in *parB*_null_ and *parS*_null_ strains, suggesting that here the ParB involvement is a part of a more complex regulatory circuit. No changes were observed in the transcription of genes adjacent to *parS5* and *parS7*-*parS10* upon ParB excess. Indeed, a recent ChIP-seq analysis did not indicate binding of ParB to these sequences *in vivo* in cells grown in minimal medium [33], but instead showed the presence of nine additional ParB-bound sites. However, none of the genes adjacent to those regions changed expression in response to ParB excess in cells grown in conditions tested here (data not shown). It is likely that the interaction of ParB with such secondary ParB binding sites, which is not required for chromosome segregation, may strongly depend on experimental conditions. It would be therefore interesting to compare ParB ChIP data obtained for cells grown in minimal vs. rich medium.

A comparison of the transcriptional changes in *parB -*deficient cells and cells expressing ParB excess identified 87 genes found in both studies (Table S4) [39]. Expression of 83 genes similarly responded to either lack or abundance of ParB. Majority of them seem to be involved in the stress adaptation. Only *PA0293*, *PA1894*, *PA2113* and *PA2792* showed opposite trends of the response. These four genes are downregulated in *parB -*deficient cells and upregulated in response to ParB excess, hence it is unlikely that, akin to genes in the proximity of *parS3* and *parS4*, ParB directly affects their expression.

Our data indicate that ParB excess downregulates two operons *pqsABCDE* and *phnAB* involved in quinolone quorum sensing signalling [68,69], probably indirectly by repressing *PA1003* (*mvfR/pqsR*) coding for a transcriptional activator of the both operons [70,71]. Additionally, the ParB excess interferes with expression of key virulence determinants, two pathways involved in the production of the siderophores pyochelin and pyoveridine.

The extreme adaptability and excellent survival of *Pseudomonas* species have been claimed to originate from the plasticity of gene expression facilitated by the high number of transcriptional regulators encoded in their genomes [83,84]. ParB excess leads to the induction of several genes encoding known and putative transcriptional regulators (Table 4). Among them is *PA2432* encoding the bistable response regulator BexR [67]. Bistability refers to a phenotypic heterogeneity within an isogenic population which allows a fraction of cells to survive in otherwise lethal conditions [85]. The BexR regulon comprises a diverse set of up- and down- regulated genes related to virulence and quorum sensing [67]. Importantly, the *bexR* induction in response to ParB excess is highly reproducible and approaches the ~10-fold induction maximum found in the subset of cells which have switched on the BexR regulon [67]. The mobilization of pKGB9 (*araBAD*p-*parB*) and pKGB8 (*araBAD*p) plasmids into the test strain PAO1161::*bexR*p-*lacZ* with “OFF” phenotype demonstrated “ON” state of *bexR*p in nearly all conjugants with pKGB9 and in a tiny fraction of conjugants with pKGB8 (<0.1%). This confirms that the ParB excess induces the *bexR* gene in all cells rather than in a specific subpopulation only. So far the mechanism of *bexR* induction in response to ParB excess remains unknown.

Our study also revealed that ParB excess leads to the induction of bacteriophage and pyocin encoding genes, which in *P. aeruginosa* are hallmarks of the SOS response [62,86]. Whereas the SOS response had initially been linked exclusively to DNA damage signals, it was later established to be part of a broad stress reaction [87,88]. SOS response is tightly connected to cell growth inhibition, cell-cycle checkpoints and even programmed cell death [89–91]. The phage- and pyocins-related genes are among the genes responding similarly to the *parB* deficiency and ParB excess (Table S4), suggesting that any imbalance in the ParA/ParB/*parS* system may trigger SOS response.

Altogether, our data indicate that the level of partitioning protein ParB in *P. aeruginosa* influences the expression of numerous genes including those involved in virulence, stress response and quorum sensing. Further studies are needed to decipher whether these effects are a direct result of ParB binding to DNA (specific or non-specific), interactions with other proteins or are caused indirectly, for example by topological changes.

## Acknowledgements

We thank dr Paulina Jęcz for the construction of pPJB132. This work was supported by the Polish National Science Centre grant no 2013/11/B/NZ2/02555 awarded to GJB.

## Supporting information captions

**S1 Figure.** Analysis of genes identified as altered in ParB+ *vs* EV but not ParB+++ *vs* EV. Abbreviation ParB+ corresponds to PAO1161 (pKGB9 *araBAD*p-*parB*) strain grown without arabinose, EV corresponds to PAO1161 (pKGB8 *araBAD*p) grown with arabinose and ParB+++ for PAO1161 (pKGB9 *araBAD*p-*parB*) cells grown with arabinose. (A) Fold change- and *p*- values for the 35 genes in ParB+ vs EV and ParB+++ vs EV analysis. First subgroup of genes with -1.6> FC >-2 or 1.6< FC <2 and *p*-value <0.05 in ParB+++ vs EV comparison is indicated in green. The second subgroup of genes with 0.05< *p*-value <0.1 in ParB+++ vs EV comparison is indicated in blue. The third subgroup of genes with *p*-value >0.1 and -1.5< FC < 1.5 is left uncoloured. Genes in operons are marked with arrows. (B) Impact of arabinose on the expression of selected genes. PAO1161 (pKGB8 *araBAD*p) cells were grown in L broth containing chloramphenicol with or without 0.02% arabinose. RT-qPCR was performed cDNA synthesized on RNA isolated from cells harvested at OD_600_ 0.5. Data represent mean ±SD from three biological replicates. Expression values for all genes are shown relative to the cells from cultures without arabinose. * - *p*-value < 0.05 in two-sided Student’s *t*-test assuming equal variance.

**S1 Table.** Oligonucleotides used in this work.

**S2 Table.** Transcriptomic changes between analyzed PAO1161 strains. (**A**) Comparison of PAO1161 (pKGB8 *araBAD*p) (EV) *vs* PAO1161 (WT). (**B**) Comparison of PAO1161 (pKGB9 *araBAD*p-*parB*) grown without arabinose (ParB+) *vs* EV. (C) Comparison of PAO1161(pKGB9 *araBAD*p-*parB*) grown with 0.02% arabinose (ParB+++) *vs* EV. (D) List of 211 loci displaying altered expression in ParB overproducing cells.

**S3 Table.** K-means clustering of ParB dependent loci. Mean levels of transcripts for all loci are shown relative to PAO1161 (pKGB8 *araBAD*p).

**S4 Table.** Genes with altered expression in response to ParB deficiency (PAO1161 *parB_null_*) and ParB overproduction [PAO1161 (pKGB9 *araBAD*p-*parB*) grown with or without 0.02% arabinose vs PAO1161 (pKGB8 *araBAD*p)].

